# Vocal plasticity in harbour seal pups

**DOI:** 10.1101/2021.02.10.430617

**Authors:** Laura Torres Borda, Yannick Jadoul, Heikki Rasilo, Anna Salazar Casals, Andrea Ravignani

## Abstract

Vocal plasticity can occur in response to environmental and biological factors, including conspecifics’ vocalisations and noise. Pinnipeds are one of the few mammalian groups capable of vocal learning, and are therefore relevant to understanding the evolution of vocal plasticity in humans and other animals. Here, we investigate the vocal plasticity of harbour seals (*Phoca vitulina*), a species with vocal learning abilities attested in adulthood but not puppyhood. To zoom into early mammalian vocal development, we tested 1-3 weeks old seal pups. We tailored noise playbacks to this species and age to induce seal pups to shift their fundamental frequency (F0), rather than adapt call amplitude or temporal characteristics. We exposed individual pups to bandpass-filtered noise, which purposely spanned – and masked – their typical range of F0s, and simultaneously recorded pups’ spontaneous calls. Seals were able to modify their vocalisations quite unlike most mammals: They lowered their F0 in response to increased noise. This modulation was punctual and adapted to the particular noise condition. In addition, higher noise induced less dispersion around the mean F0, suggesting that pups may have been actively focusing their phonatory efforts to target lower frequencies. Noise masking did not seem to affect call amplitude. However, one seal showed two characteristics of the Lombard effect known for human speech in noise: significant increase in call amplitude and flattening of spectral tilt. Our relatively low noise levels may have favoured F0 shifts while inhibiting amplitude adjustments. This lowering of F0 is quite unusual, as other animals commonly display no F0 shift independently of noise amplitude. Our data represents a relatively rare case in mammalian neonates, and may have implications for the evolution of vocal plasticity across species, including humans.

## INTRODUCTION

### Animal communication and plasticity

Communication involves the transmission of information between two or more individuals [1]. Accurate communication can increase an animal’s fitness; therefore, it seems essential to understand the possible effect of biotic and abiotic factors on communication. In fact, communication can increase e.g. potential mating opportunities, the probability of escaping from a predator, and the speed of social learning [2]. Acoustic communication is particularly well-developed in marine mammals because of the selection pressures of the marine environment [3, 4]. Underwater sounds propagate over long distances whereas water clarity or light level can limit transmission of chemical or visual cues [5]. Signallers can face unwanted noise that leads to signal degradation. Masking occurs when the frequencies of the noise overlap with the frequency range of the signal [6].

The current study targeted one building block of animal communication systems: vocal plasticity. Being vocally plastic allows individuals to adjust their vocal signals in response to changes in their environment [6]. Plasticity, if present in a species, can be deployed in various contexts ranging from environmental challenges to signal detection to changes in social interactions. Interferences in signal detection can lead to important adaptations underlying the evolution of animal communication systems. Vocal plasticity enables some animals, including humans, to reach their communicative goal, potentially via different mechanisms.

One type of signal modification requires little plasticity and is common across species. In particular, many birds and terrestrial mammals increase amplitude levels of vocalisation in the presence of masking noise [7–10], especially when the noise overlaps with the spectral composition of species-typical vocalisation [11, 12]. This adjustment of vocalisation amplitude in response to background noise is called the Lombard effect [13], which has been well-studied in humans.

### The Lombard effect

When it comes to human communication, the Lombard effect is common and more prominent when speaking occurs with communicative intent with a speaking partner than when speaking aloud alone [14]. When speaking without communicative intent, one function of the Lombard effect may be to monitor the speaker’s own speech for errors [15]. The type of voice modification and its strength varies between individuals [16]. Flattening of the spectral tilt has been related to a significant increase of intelligibility of speech under noise. Conversely, an increase in F0 has not shown a significant increase in intelligibility [17], and thus might be a side-effect of increased subglottal pressure to achieve a higher intensity [18, 19]. Human speakers start showing the Lombard effect when the background noise reaches 40-50 dB of sound pressure level (SPL). In particular, the Lombard effect starts appearing at 43.3 dB SPL of pink background noise, after which the SPL of the speaker increases by 0.65 dB per 1 dB of added noise level [20].

Recent animal evidence shows that the signal-to-noise ratio (SNR) between animal vocalisations and background noise is a better predictor for the Lombard effect than the ambient noise level alone. Lack of Lombard effect in experiments may thus be due to too low SNR, or species-specific characteristics, such as always vocalising close to the physical limits, even in the absence of noise [21, 22].

Though amplitude adjustment is only one, extremely common, adaptive strategy. When flexibly adapting their vocal output, some species exhibit rare spectral changes [23–26], while others show different vocal behaviours, such as temporal shifts which might include, among others, changes in call rate or duration [27–29].

### Spectral adjustments in animal communication

Parallel strands of research investigate vocal plasticity and vocal production learning, which can also arise from plastic adaptations to environmental factors, but complex forms involve modulation of F0 or formants. Vocal learning is the capacity to produce novel vocalisations through imitation or experience. This ability is often, but not always, supported by control of vocal articulators and cavities (i.e., the manipulation of the vocal folds, larynx and supralaryngeal vocal tract [30]).

Beyond vocal learning, other forms of vocal parameters modifications can reflect dynamic and static attributes such as the physical, motivational or arousal state of an organism. For instance, vocalisations produced by larger species or larger animals within a species are usually characterized by lower formant frequencies [31]. Adjustment of acoustic parameters in vocalisations often occurs via permanent anatomical structures and contexts involving emotional arousal. Therefore, many cases do not require volitional modifications of vocal parameters and require little to no plasticity. For instance, some species possess unique anatomical features and produce unusually low formant spacing, such as koalas and deers [32, 33]. Several species show variability in their F0: vocalisations emitted by domestic dogs (*Canis lupus familiaris)* and wapitis (*Cervus elaphus*) in an aggressive context have a lower F0 than vocalisations emitted in a positively valenced context [34–37]. Other species, such as baboons (*Papio hamadrayas ursinus)* [38] or bats (*Megaderma lyra)* [39], show an increase of their F0 during agonistic or stressful interactions. Perceptually, frequency parameters can affect communication of physical and emotional characteristics: receivers may rely on vocal characteristics to gauge information about the emitter and may adjust their own behaviour accordingly [40, 41]. To summarize, anatomical adaptations and emotional contexts can affect a species’ F0 with no need for plasticity or control.

In contrast, the volitional and controlled modulation of vocal parameters, such as F0, are rarely observed in mammals. Few studies have demonstrated the capacity of fine F0 modulation [42, 43], sometimes coinciding with vocal imitation (elephants: [44, 45]). Some bats are capable of vocal imitation [46] and pale spear-nosed bats (*Phyllostomus discolor)* can be trained to shift their F0 downwards [47, 48]. Birds increase their F0s in urban environments due to low frequency traffic noise [49]. However, F0 shifts due to noise are considered rare in animals and, when they occur, may be driven by the Lombard effect as a physiological by-product of higher vocal amplitude [50].

### Our approach

Methodologically, plasticity in animal vocal behaviours can be investigated in several ways. Studies in the laboratory provide the advantages of an experimentally controlled set up, but also the challenge to obtain spontaneous vocalisations [12, 51, 52]. In contrast, the use of operant conditioning techniques has proven effective in eliciting vocalisations. If production of vocalisations involves operant learning, it can be difficult to disentangle natural predispositions towards a task from learning attitudes [12, 26, 29, 53]. At the other extreme, fieldwork favours naturalness and spontaneity, sometimes at the expense of experimental control. Here, we attempted to combine the best of these empirical approaches. First, we tested wild animals soon after they reached captivity. Second, we capitalized on their natural proclivity to spontaneously produce vocalizations. Third, we played noise that had been recorded in their natural environment, but filtered it to target a specific frequency range to test our specific hypotheses.

Previous studies targeting amplitude shifts in presence of background noise of marine mammals have mainly focused on cetaceans [23–25, 27, 54]. In harbour seals, adult males did not increase the amplitude of their vocalisations in response to vessel noise [55]. Had F0 been investigated, it would have probably also been unaffected by noise, because the noise did not fully overlap with the frequency range of adult seals’ displays. In contrast, we decided to investigate the plasticity of F0 in seal pups. Understanding harbour seals’ vocal plasticity is important for three main reasons. First, harbour seals face ecological challenges which may impact communication in their natural environment. This species forms unstable seasonal groups during pupping season, with up to several hundred females sharing an area for the entire lactation period [56]. These environmental conditions provide ambient noise which is variable in amplitude and affected by factors such as weather disturbances and anthropogenic noise. For instance, wind farms installed around the world emit a relatively weak but characteristic noise (usually around 50 dB). Second, harbour seals are highly vocal during puppyhood. They have a vocal recognition system enabling mate attraction, mother-pup recognition, and contact maintenance [57]. Females, just a few days postpartum, alternate between foraging trips to sea and nursing their pups on land [58]. Thus, pups may be separated from their mothers either on land or at sea, and successful reunions also depend on the characteristics of mother attraction calls (henceforth “calls”) emitted by pups [57, 59]. Third, some adult pinnipeds are capable of vocal production learning [26, 60, 61] which is the ability, rare among mammals, to modify species-specific vocalisations or create novel ones, often through imitation [43]. Vocal plasticity in harbour seal pups can provide a developmental window into mammalian vocal learning. Among mammals, pinnipeds are an excellent model and one of the few clades showing vocal learning; they are phylogenetically closer to humans than other classical models for vocal learning (e.g. songbirds) and exhibit a variety of spontaneously produced vocalisations [60, 62, 63]. Finally, vocal plasticity in pinniped puppyhood could also provide indirect evidence for specialized neuro-anatomical structures subserving vocal learning.

### Hypotheses and predictions

In the current study, we aimed at triggering punctual shifts in F0 and other vocal parameters, in a controlled experimental setting devoid as much as possible of emotional features. Our goal was to induce vocal shifts which would be volitionally produced as a strategy to avoid acoustic masking in a noisy environment, thus illustrating unusual vocal plasticity in this promising taxon.

We predicted that, to compensate for a noisy environment, pups would increase signal amplitude or shift vocalisations temporally or spectrally. Thus, we should observe some changes of their vocal parameters (e.g. amplitude, F0, duration) in their calls according to different amplitudes of background noise.

The first and main aim of this experiment was to induce a shift in the seals’ F0. Thus, we expected pups to shift their F0 upwards or downwards to escape the bandpass-filtered noise which we purposely tuned to overlap with their F0 range. Alternatively, a lack of F0 shift could support hypotheses of less reliance on F0 adaptations in social communication, or lack of the vocal plasticity necessary to conduct such modulations. Considering pinnipeds’ vocal production learning capacities in adulthood, we favoured the former hypothesis.

The second aim of this experiment was to test for two additional types of modulation. If seals behaved similarly to other species, we would also expect temporal shifts in the pups’ vocalisations and in their numerosity [28, 64–66]. In particular, seals may modulate the rate and duration of calls to maximize information transmission during noise. If pups performed such adjustments, we would expect more and longer vocalisations during noisy periods when compared to the absence of playback [28]. While we did not expect a difference in number of calls, we hypothesized we might find temporal adjustments [29].

Finally, if the Lombard effect seen in other species also applied to harbour seals, we would expect pups to exhibit an increase in their vocalisations’ amplitude during playbacks of lower amplitude noise compared with no playbacks, and even more so during playbacks of higher amplitude noise. Conversely, no amplitude shifts in the vocalisations would suggest that higher noise levels may be needed. Alternatively, lack of amplitude adjustments could be due to seals adopting a different (spectral or temporal) strategy in response to masking. Based on evidence in adult harbour seals, we favoured the hypothesis of no amplitude adjustment [55].

Overall, there could be a trade-off between vocal adjustments, leading to an amplitude modification, an F0 variation, or temporal changes. In particular, the combined outcome of frequency and amplitude shift could shed light on neurobiological and biophysical mechanisms. While a lack of both frequency and amplitude shifts would confirm the results obtained in adult seals [55], an amplitude-only shift would point towards a general Lombard mechanism, which is broadly conserved across species. A simultaneous shift in F0 and amplitude could suggest that the F0 shift was a mechanical by-product of the amplitude shift. Finally, a frequency-only shift may point towards vocal plasticity not as a by-product of Lombard modulation, but possibly due to good neural control of the larynx.

## MATERIAL AND METHODS

### Subjects and study site

The study was conducted at the Sealcentre Pieterburen, a seal rehabilitation centre specialised in phocids. The Sealcentre rescues a yearly average of 400 seals (family Phocidae), later released back into the wild. Tested individuals were housed in quarantine units. Data collection started immediately after arrival of the individuals and once a veterinarian confirmed animals were not suffering from any extenuating disease (Table S1 in Supplement). To maintain the animals as wild as possible, seals were in contact with humans only during the four daily feedings, which occurred independently from the experiment. All recordings described here were performed with no water in the pool, hence avoiding water noises.

Subjects were wild-born, Eastern-Atlantic harbour seal pups. This species is monotocous, granting that animals could not be siblings. Expert veterinarians estimated the pups’ age during the first veterinary examination following the Sealcentre’s protocols [67]. Data was collected from 8 seal pups (3 females), aged between 7 and 10 days on the first day of testing (June 25, 2019). Seals were housed in pairs, each pair in a separate quarantine. Housing conditions of all four quarantines were identical.

Data collection was non-invasive, approved by the centre’s veterinarians, and adhered to the guidelines of the Association for the Study of Animal Behavior [68]. We observed that the noise playback did not increase the pups’ behavioural indicators of stress by live video monitoring the first playbacks under the supervision of the research and veterinary team.

### Stimuli

Playbacks were based on audio recordings of ambient noise from a sandbank in the Wadden Sea (see Supplement). Sounds were bandpass filtered in Praat (version 6.0.52; [69]), resulting in a noise band between 250 and 500 Hz. This frequency range was chosen to overlap with the F0 range of seal pups’ mother attraction calls [57, 70, 71].

An experimental playback session consisted of a 45-minutes audio file (WAVE format) composed of three sequences of 5-minutes *high noise* (65 dB SPL), three sequences of 5-minutes *low noise* (45 dB SPL), and three sequences of 5 minutes with *no playback* (resulting in approx. 25 dB SPL of background noise). Prior to experimental trials, the playback noise was measured with a SPL meter at the centre of the dry pool at a seal pup’s height (approximately 30 cm).

The order of sequences within playbacks was randomised and constrained by avoiding two identical amplitude conditions in a row (e.g. 2 high-noise playbacks in a row). This was aimed at preventing the pups’ habituation and increasing the chances of observing a punctual noise-induced voice modulation. This combinatorial constraint resulted in sixteen playbacks, that were the only and all possible ordered combinations of the previous sequences described (no playback, low noise, and high noise) each appearing three times. A playback was then randomly selected for every seal pair and experimental session.

### Apparatus and experimental procedure

Sounds were played via a Yamaha HS5 Speaker (2-way bass-reflex bi-amplified nearfield studio monitor; 38 Hz–30 kHz (−10 dB), 47 Hz–24 kHz (−3 dB) frequency response). All recordings were performed with a unidirectional microphone Sennheiser ME-66 (frequency response: 40 Hz–20 kHz; Sennheiser electronic GmbH & Co. KG, Wedemark, Germany) on a tripod. During playback, this microphone, connected to a Zoom F8 recorder (Zoom Corporation, Tokyo, Japan), recorded the pups’ vocal responses. The apparatus was positioned at approximately 2 meters from the pup at one corner of the pool (Figure S1 in Supplement).

We tested four pups per day (one session a day in two quarantines). Once we obtained 7 valid sessions (containing at least 2 vocalisations), we carried on the experiment on four additional pups. The first quarantine was tested at 2:15 p.m. and the second one at 6:15 p.m. These times were chosen, in agreement with the Sealcentre’s veterinarians, to increase the likelihood to successfully record spontaneous vocalisations because pups are usually more vocal before feeding. The apparatus was temporarily installed in each quarantine three hours before each session (i.e., 11 a.m. and 3 p.m.) and directly removed after. By the end of the study, all animals had been recorded between 10 and 14 days to reach 7 valid sessions.

### Sound recordings, annotations, and F0 extraction

Acoustic analyses were carried out in Python and MATLAB once the recorded files (WAVE format) were manually annotated in Praat 6.0.52 [69]. The annotation process, which was manually performed twice, consisted in annotating the onset and offset of each vocalisation (Praat settings: view range = 0-3000 Hz, window of analysis = 0.05 s, dynamic range = 70 dB).

A Zoom Q8 handy video recorder filmed every trial. For each quarantine, the video was analysed using BORIS [72]. As two individuals were housed in the same quarantine, one was marked with an animal waterproof coloured marker. Audio and video recordings were synchronized to assign each vocalisation to the pup that produced it.

After annotation, the Parselmouth Python library (version 0.3.3, Praat version 6.0.37; [73]) was used to extract duration and F0 of the annotated calls (autocorrelation method for pitch tracking, with non-default parameters: time step 0.01 s, pitch floor 200 Hz, and pitch ceiling 800 Hz). All calls were included in the analyses of the number of calls and their duration. However, only calls which 1) were not clipped, 2) did not overlap with other individuals, 3) did not contain background noise other than the playback, and 4) could be properly tracked by Praat were included in the analyses of calls’ amplitude and F0.

Praat’s ability to track the pitch in all noise conditions was checked manually by two researchers on a large random sample of calls. This was first done by zooming in on the sound wave, selecting a single period, and calculating the frequency as the inverse of length of that period. Secondly, the spectrograms were visually verified, checking whether estimates by Praat matched the F0 and harmonics in the spectrogram. By doing so, we did not find any bias due to the pitch tracking algorithm’s performance in our recordings: even in cases where high intensity noise condition obscured the F0 in the 250-500 Hz frequency band, the harmonics provided enough autocorrelation information for Praat’s algorithm to estimate F0.

### Amplitude and spectral tilt

We obtained average spectra and intensity values for each call to test if seals adjusted their vocalisations’ amplitude or spectral tilt depending on the noise condition. To account for the differential contribution of noise in each condition, separate recordings were made of noise only (seals not present) with otherwise equal recording setup. The intensity and the spectral characteristics of the background noise was seen to vary slightly over time, and noise-only recordings allowed for more accurate estimation of the noise characteristics during each vocalisation. Due to the very reverberant recording conditions and additional noise sources (e.g., bird and airplane sounds), perfect cancellation of the playback noise from each recording was not possible. For comparisons between conditions, we tried to reduce the effect of the noise based on spectral subtraction, by subtracting the averaged power spectrum of the estimated background noise from that of the vocalisation [74]. Background noise increased the mean and the variance of the spectral content of the underlying calls. Spectral subtraction can recover the mean spectral content, but the variance will remain distorted by the noise variance [74].

Each recording session had slightly varying preamplifier gain in the recording phase due to manual adjustment. This gain variation was compensated by 1) calculating the root-mean-square (RMS) power for each noise condition from each recording from the moments when the seals were not vocalizing and 2) determining a gain value per recording session, that brought the average power of the low and high noise conditions to the same level as the corresponding average value in the noise-only recordings.

To perform call amplitude analysis, we calculated the RMS power of each call (RMS power is proportional to the RMS sound intensity, and for simplicity, this measure will be called *intensity* from here on). Similarly, we calculated the intensity of the noise-only recording from the corresponding location and subtracted it from the intensity of the call. If the signal-to-noise ratio of a call was too low, this subtraction could lead to a negative intensity value for the call. However, as the mean over all calls after the subtraction should represent the mean of the original calls, comparisons in the linear (non-decibel) domain were possible.

The Lombard effect on human speakers shows as an energy boost on high frequencies. This can be characterised by, for example, a flattening of the spectral tilt. In [17, 75] speech in noisy conditions showed as a spectral energy boost between 0.5–1 kHz and 5 kHz when compared to the silent condition. Different ways to measure spectral tilt show high variance [76], thus, in the current study, spectral tilt was estimated using two separate methods.

First, after spectral subtraction, spectral slope was calculated by fitting a regression line in the log-energies on ⅓ octave frequencies as done in [17]. In this work, 1 kHz was used as a reference frequency for the ⅓ octave filters, and the line was fitted only on frequencies above 400 Hz. We adopted this cut-off because the F0 of the vocalisations lied around this frequency, and the background noise corrupted mostly estimates of the spectral energy under 500 Hz. The regression line was fitted only on the octave energies whose values remained positive after spectral subtraction. For 11 calls, octave slope could not be estimated due to lack of positive energy value on two or more bins after spectral subtraction. These occurred only in the high-noise condition, and were discarded from the spectral slope analysis.

Second, again after spectral subtraction, a ratio of the spectral energy between 0.4 and 1 kHz to that of 1–4 kHz was calculated (R14). This method was adjusted from [77]: instead of considering all energy below 1 kHz as in their work, we removed energies below 400 Hz from the analysis as justified above. For spectral analysis for each call and corresponding noise estimate from the noise-only recording, the average power spectra were calculated using FFT with window size of 512 samples, overlap of 256 samples and Hamming windowing.

### Statistical analysis

The effect of the noise intensity on the number of vocalisations, call duration, and F0 was analysed by fitting a linear mixed-effects model. These extracted acoustic parameters were included as dependent variables, predicted by the background noise condition as an independent variable (factor with three levels: low noise, high noise, and no playback). The session number (7 sessions per pair) and the specific seal identity were modelled as random effects. We also included a variable named “trial number” to control the existence of a learning or habituation effect within sessions. This variable allowed us to test whether changes in vocal behaviour were affected by the time course of the session.

Statistical analyses were performed in R, version 3.5.2 [78]. Comparisons were done with linear mixed-effects models (LMM) using the package `nlme` [79]. P-values were calculated via Monte-Carlo sampling with 1000 permutations using the PermTest function of the R package `pgirmess` [80]. Permutation tests for linear models were chosen because suited our limited sample size and relaxed the assumption of normality of residuals [81]. Moreover, a Bonferroni adjustment for multiple comparisons was applied to all pairwise comparisons. Significance was set at *p* < 0.05 / 3≈ 0.0167.

To analyse the effect of the intensity of playback noise on the intensity of the seals’ vocalisations and the two spectral tilt measures described above, we used the non-parametric Mann-Whitney U test. These three variables were analysed differently to the previous ones, since they were strongly non-normally distributed and as such could not be fitted by linear models. To control for the individual effect of seals, tests were done per seal. To correct for multiple comparisons, we applied a Bonferroni correction of 24 (8 seals, 3 pairwise comparisons between the three noise conditions), resulting in a required significance level of *p* < 0.05 / 24 (≈ 0.00208).

## RESULTS

We recorded a total of 3534 calls. We tested 8 pups and obtained 7 valid sessions per pair (mean: 12 days, min: 10 days, max: 13 days). Statistical analyses conducted on vocalisations’ amplitude and F0 were performed over 2576 ‘clean’ calls (see 4 criteria in Methods). Statistical analyses on vocalisations’ rate and duration were performed over the totality of recorded calls.

### Number of calls

There was no significant effect of the noise condition on the number of vocalisations (pseudo*R*^*2*^ = 0.027; *p* = 0.341; *N* = 245). Thus, pups did not significantly increase or decrease their number of vocalisations depending on the noise amplitude (Figure 1). We recorded in total 1209 calls in high noise, 1227 calls in low noise and 1097 calls in no playback. The number of vocalisations was also comparable throughout the conditions.

**Figure 1.**
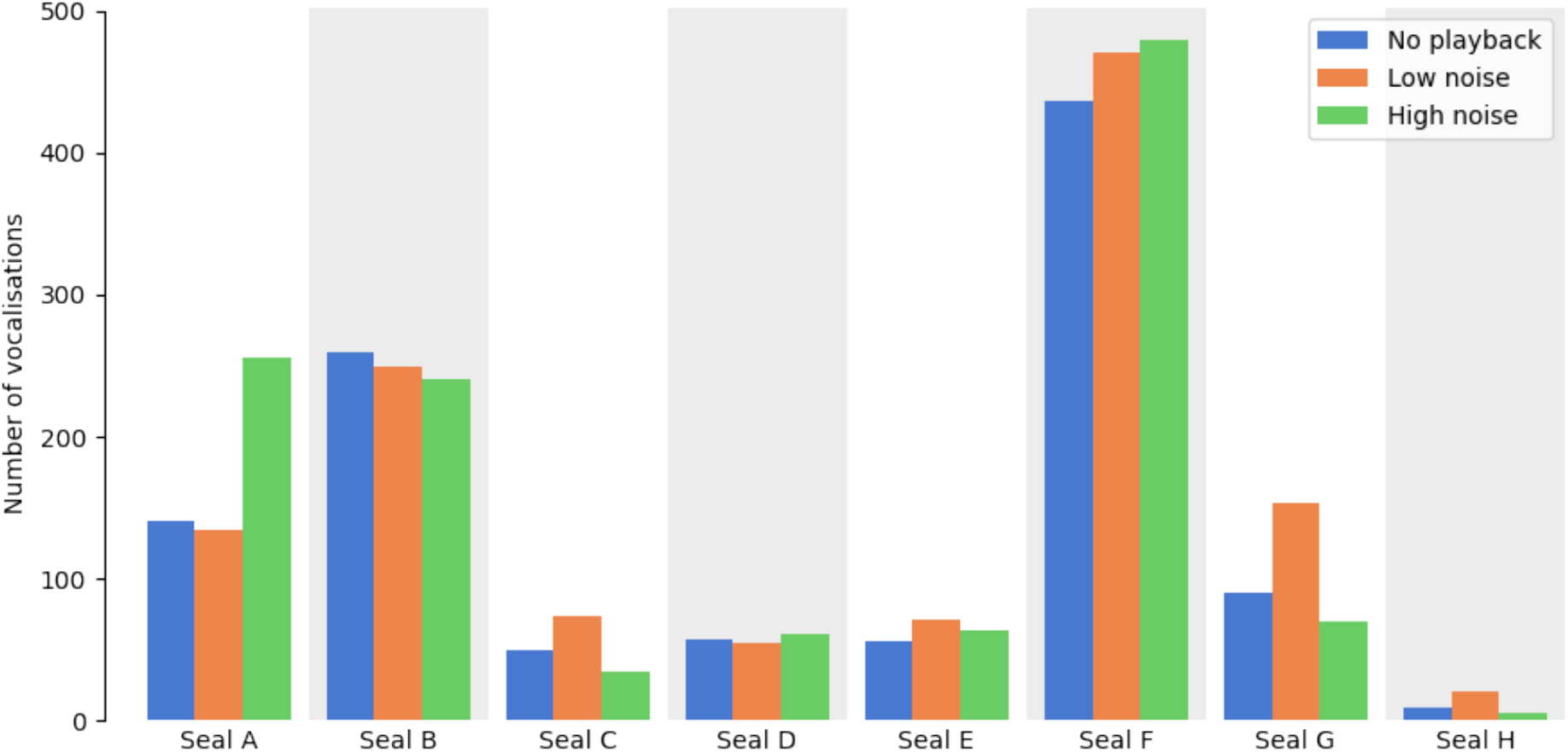
Amount of calls per seal grouped by condition. There was some variation in number of recorded vocalisations among conditions and seals, but there was no consistent effect between conditions.

### Duration

Noise conditions did not affect calls’ duration (Figure 2). Vocalisations were neither significantly longer nor shorter as noise level increased (pseudo*R*^*2*^ = 0.014; *p* = 0.707; *N* = 3534). Pups’ calls lasted 0.785s on average (median: 0.729 s; min: 0.182 s; max: 3.892 s).

**Figure 2.**
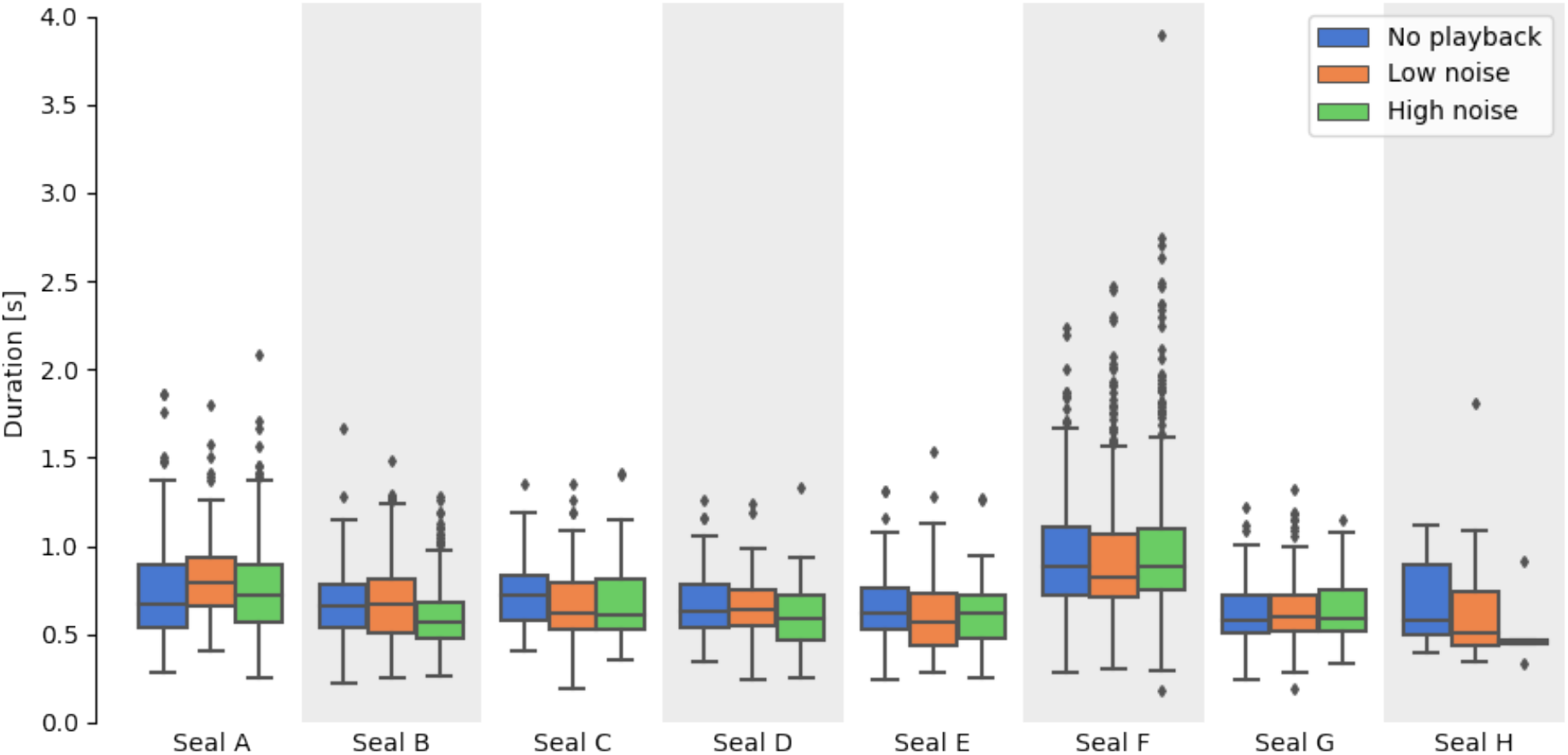
Duration of vocalisations. There was no significant effect of the three conditions on the duration.

### Fundamental frequency (F0)

We tested the effect of the noise condition on F0 (Figure 3). A main significant effect was found (pseudo*R*^*2*^ = 0.202; *p* < 0.001; *N* = 2576). Pairwise comparisons showed also significant differences between our three levels. In high noise, F0 was significantly lower than in low noise (pseudo*R*^*2*^ = 0.166; *p* < 0.001; *N* = 1751) and no playback (pseudo*R*^*2*^ = 0.287; *p* < 0.001; *N* = 1636). F0 was also significantly lower in low noise than in no playback (pseudo*R*^*2*^ = 0.038; *p* < 0.001; *N* = 1765). The F0 median in the high noise condition was equal to 324 Hz, 374 Hz in the low noise condition, and 403 Hz in the no playback condition.

**Figure 3.**
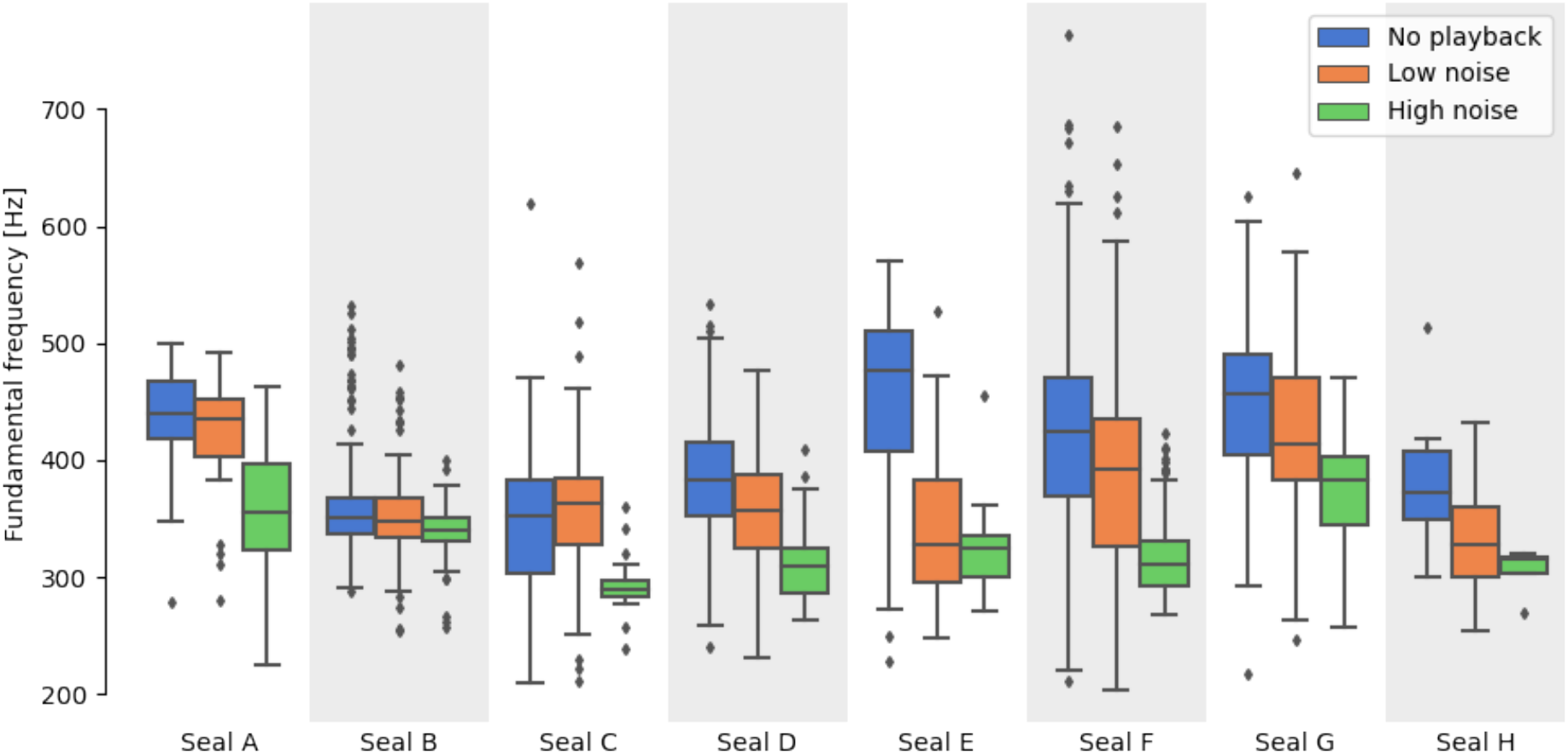
F0 of the seal’s vocalisations. The three different conditions of noise intensity had a significant effect on F0.

We tested for the existence of a learning or habituation effect within sessions. Because of our playback duration, we could have observed differences in vocal behaviour between the beginning and the end of each session due to habituation, frustration, or tiredness. We did not find any significant effect of trials on F0 (*p* = 0.184; *N* = 2576).

### Amplitude and spectral tilt

Initial Mann-Whitney U tests on the whole dataset showed no significant effects on call amplitude (see also Figure 4 and Table S2 in Supplement). After Bonferroni-correction for two measures, both spectral tilt measures showed an effect only between no playback and low noise condition (no playback vs. low noise: R14: *p* = 0.0041, slope: *p* < 0.001; no playback vs. high noise: R14: *p* = 0.025, slope: *p* = 0.47). After analyzing calling patterns of individual seals, one seal appeared to contribute most to the seen global effect (Figure 5). Follow-up analyses were performed for each seal separately.

**Figure 4.**
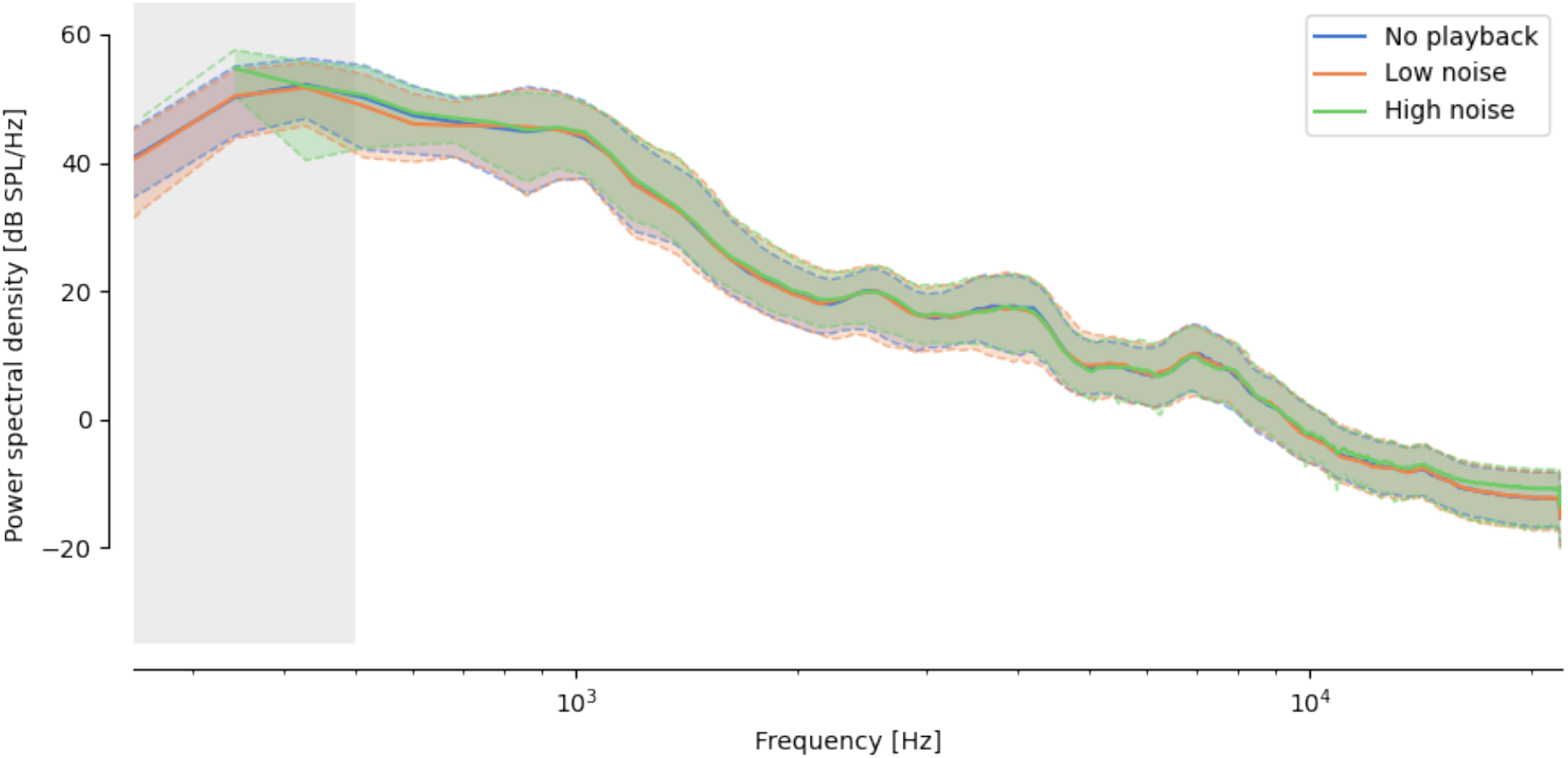
The median spectrum over all seal vocalisations grouped per noise intensity level and its first and third quartile illustrated the lack of general effect of the noise on the seals’ vocalisations. (Blue: no playback, orange: low noise, green: high noise.) See also Figure S2 in Supplement for individual spectra.

**Figure 5.**
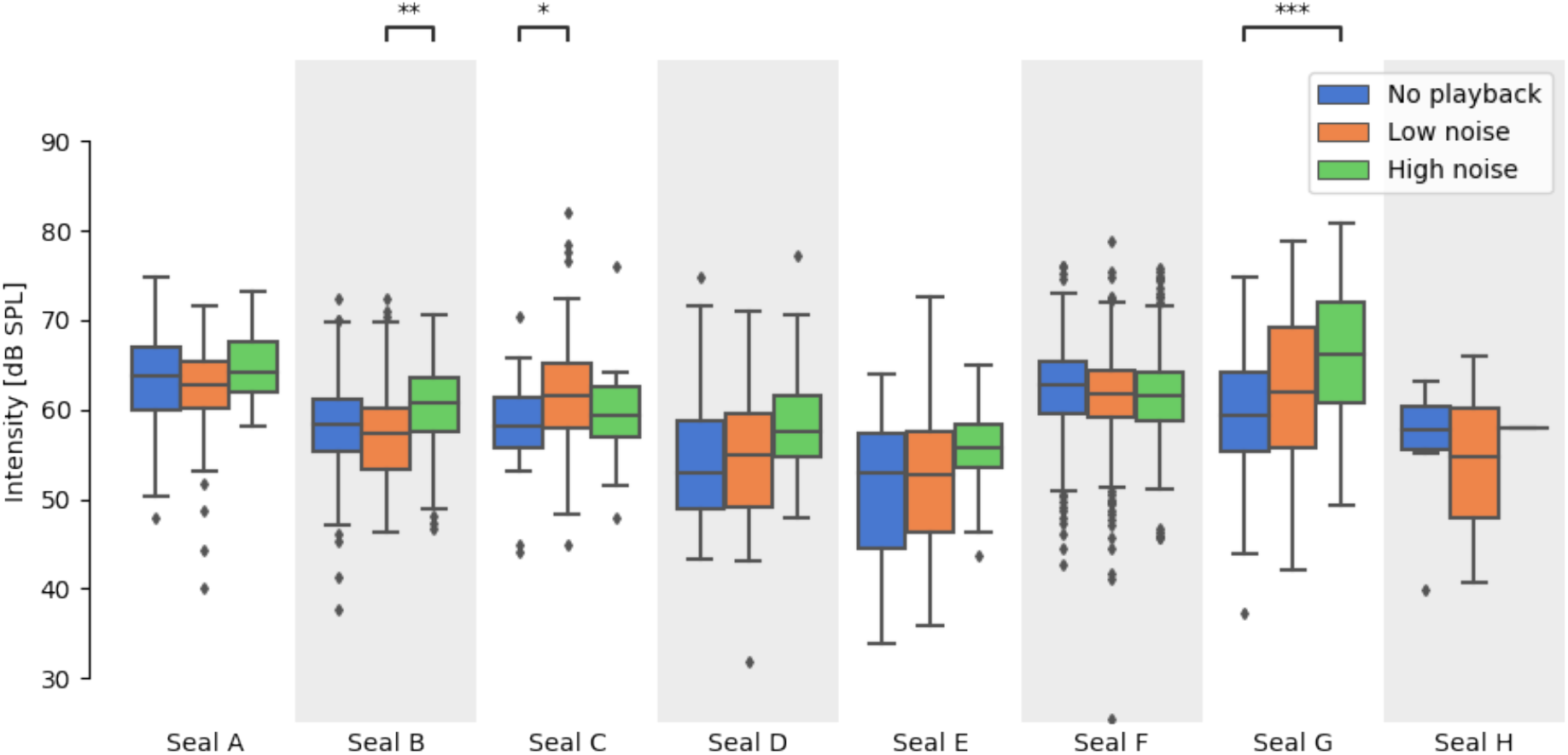
Intensity of vocalisations after compensating for the noise intensity. While there was no overall effect of the noise intensity on the intensity of the seals’ vocalisations, seal B, seal C, and in particular seal G showed a significant increase in their vocalisations’ intensity between at least two conditions. Significant differences between conditions are marked with asterisks (*: *p* < 0.05; **: *p* < 0.01; ***: *p* < 0.001; p-values are Bonferroni-corrected by factor 24).

Seal G showed increased call amplitudes with increasing noise level. The effect between no playback and high noise condition was significant (*p* < 0.001). The spectral slope was seen to flatten by 0.31 dB/octave from no playback to low noise (*p* < 0.001; Figure 6). Similarly, the spectral ratio R14 (Figure S3 in Supplement) decreased between no playback to low noise (*p* < 0.001) and no playback to high noise (*p* = 0.0011). Seal C showed significant effect in amplitude between no playback and low condition (*p* = 0.0012), but no other significant effects following the Lombard hypothesis. Seal B showed a significant effect in intensity (*p <* 0.001) and R14 (*p* = 0.0019) only between the low noise and high noise conditions.

**Figure 6.**
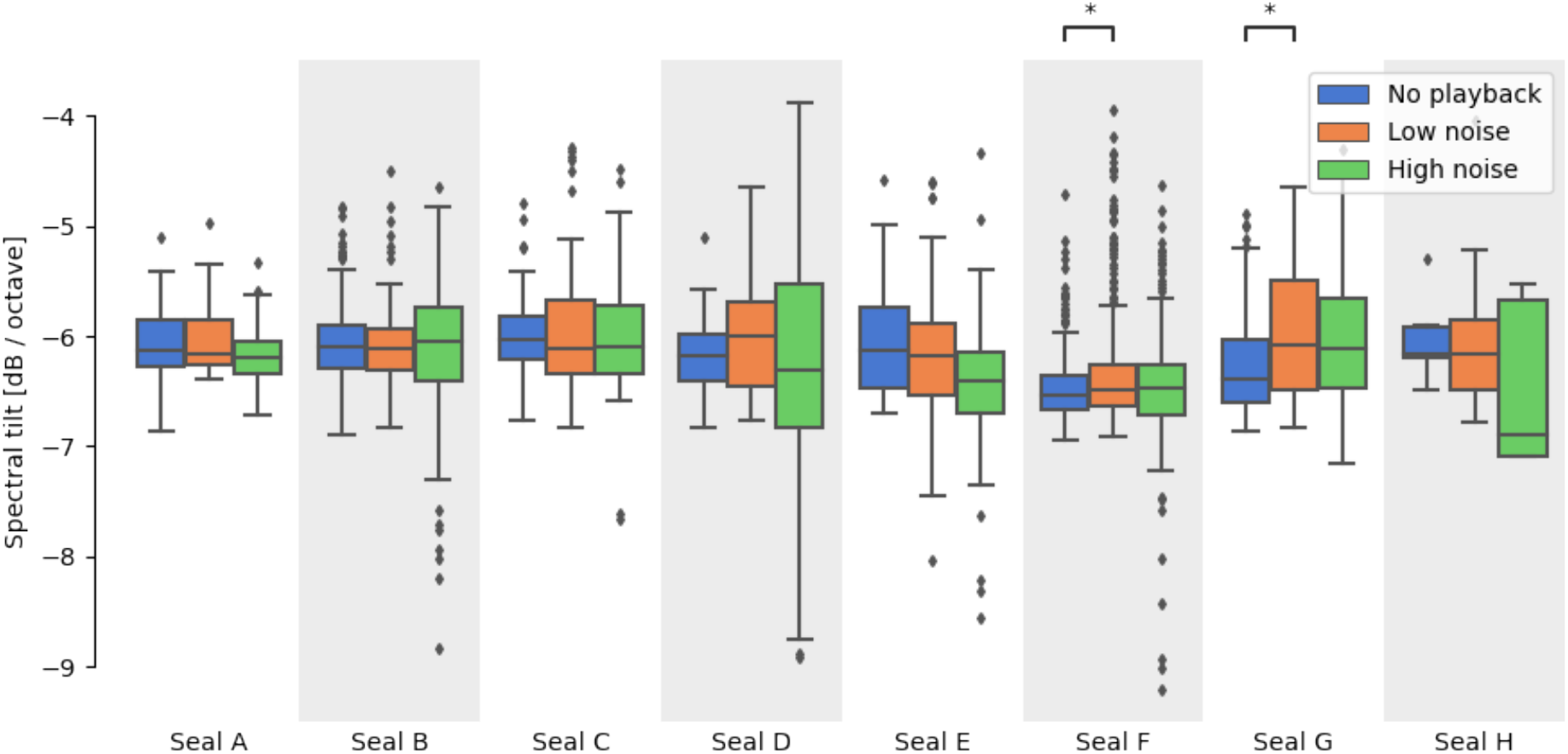
Fitted slopes of spectral tilts. The fitted slopes of the spectral tilts of seal F and seal G’s average vocalisation spectra show a flatter spectral tilt in noisier conditions and provide suggestive indication of the Lombard effect potentially occurring in these individuals. Significant differences between conditions are marked with asterisks (*: p < 0.05; **: p < 0.01; ***: p < 0.001; p-values are Bonferroni-corrected by factor 24).

### Coefficient of variation

Statistical analyses on the level of dispersion around the mean were conducted for vocalisalisations’ duration and F0 (Figure 3). We calculated the coefficient of variation of vocalisations grouped by session, seal identity, and condition. No significant differences between conditions were found on the coefficient of variation of calls’ duration (pseudo*R*^*2*^ = 0.002; *p* = 0.886; *N* = 118). However, we found a significant effect of noise conditions on the coefficient of variation of F0 (pseudo*R*^*2*^ = 0.195; *p* < 0.001; *N* = 109). Pairwise comparisons showed that the coefficient of variation of calls was significantly lower in the high noise condition compared to both, low noise (pseudo*R*^*2*^ = 0.256; *p* < 0.001; *N* = 69) and no playback conditions (pseudo*R*^*2*^ = 0.233; *p* < 0.001; *N* = 71). No significant difference was found between the low noise and no playback conditions (pseudo*R*^*2*^ = 0.004; *p* = 0.510; *N* = 78).

## DISCUSSION

### Overview of findings

Our data showed a clear downward F0 shift in harbour seal pups in response to noise masking. The number of calls and their duration were neither influenced by the presence of ambient noise nor by its amplitude levels. In addition, three out of eight pups showed limited modulation of their call amplitude depending on the noise condition, perhaps indicating compensation for acoustic masking.

Overall, we found no strong modulation of spectral tilt or call amplitude as a response to increased noise levels. Both of these quantities are usually measured when testing for the Lombard effect in human speech and animal vocalisations. Our findings are in line with previous results from adult harbour seals [55], where no effect on call amplitude was observed.

In our study, signal-to-noise ratio (SNR) in the high noise condition was approximately +5 dB on average (a noise level of 65 dB corresponded to a mean intensity of calls of 59.7 dB in no playback condition, and 60.3 dB high condition after spectral subtraction). For comparison, SNR’s of −5 to −20 dB induced the Lombard effect in frogs, whereas an SNR of +20 dB did not [21]. The underwater vocalisations of harbour seals in [55] had very high SNR (50-70 dB), perhaps also contributing to the lack of observed Lombard effect. [75] reported how the Lombard effect on human speakers helped maintain a +12.5 dB signal-to-noise ratio under noise conditions on average, where the SNR would otherwise be negative without any vocal intensity modification. Drawing on this evidence, the relatively low noise levels of our playback may not have masked seal pup vocalisations sufficiently to induce a general amplitude shift that could compensate for increasing levels of ambient noise.

### Amplitude modulation in one pup

One of the seal pups showed a peculiar vocal behaviour. Vocalisations of pup G showed flattening of the spectral tilt when background noise was present and increasing intensity as noise increased. Energy of the vocalisations on the 1–4 kHz spectral band increased more than that of below 1 kHz in response to noise, similarly to the Lombard effect in human speakers. The flattening in the spectral tilt observed for pup G under noise was approximately 0.31 dB/octave (no additional flattening from low to high noise condition). A similar change (flattening of 0.27 dB/octave) in spectral tilt occurs in human speakers speaking in quiet vs. 82 dB SPL background noise [17]. Our average +5 dB SNR in the high noise condition may be close to the threshold where the Lombard effect begins to take place. In addition to the strong evidence for seal pup G, this may also explain the sporadic effects of spectral tilt and amplitude modulation for pups B, C and F.

The noise threshold inducing the Lombard effect is variable between individuals [16]. Because of this, pup G may have been more responsive to noise compared to the other individuals. Furthermore, its vocal behaviour could illustrate a higher motivation and arousal induced by the noise context. Finally, and more speculatively, the increased amplitude in pup G may have arisen from stronger communicative intent compared to the other pups, as seen in humans in the context of social communication [14]. Based on the vocal behaviour of pup G, we cannot exclude that harbour seals can increase the amplitude of their voices in response to noise.

### Fundamental frequency shift

Our experimental playback successfully spurred the seals to modify their laryngeal phonation. This behaviour may be an adaptation to avoid spectral masking of one’s F0. This vocal modification was punctual and adapted to the particular noise broadcasted during this experiment. Our results show that seal pups modified their vocalisations in a unique way: a downwards F0 shift was observed in response to increased ambient noise. The lowering of F0 is atypical when compared to other species that have shown either no shift or an increase in their F0 [6, 47, 50]. Analyses on the dispersion of F0s around their means across vocalisations revealed that dispersion was lower in the high noise condition than the low noise condition and no playback condition. This suggests that, in addition to shifting down their F0, seal pups may have *focussed* their vocal production towards these lower frequencies. This downward shift of F0 could have at least two functional explanations. First, it may be an adaptation to the actual environmental noise pups encounter: as lower frequencies propagate better in wind, shifting F0 downwards may increase the travel distance of calls [82]. Second, lowering of the F0 may be a way for seal pups to better communicate their identity. As low F0s induce closely spaced harmonics, hence more frequencies per frequency band, the upper vocal tract acting as filter has a ‘denser’ source to create formants on. Indeed, close spacing of harmonics contributes to enhanced formant information [83, 84] which may be a key parameter for individuality encoding [85, 86].

The shift in F0 cannot be explained by automatic adaptations (as opposed to some vocal control). Indeed, arousal can lead to tension of the vocal folds, inducing an increase in vibration frequency and producing, in turn, an increase in F0 [87]. The downwards F0 shift we find therefore contrasts with predictions of arousal-driven F0 shifts. Indeed, our evidence in a novel context in harbour seal pups may be interpreted as a behavioural proxy for developed laryngeal control.

### Vocal plasticity and neuro-anatomical mechanisms

The present experiment setting was not designed to induce imitative behaviour or a specific vocalisation change (for example by rewarding a specific vocal parameter shift [26, 47, 48]). Instead, our playback triggered a spontaneous and volitional vocal response. Over a short time period, seal pups lowered and increased significantly their F0 according to the playback noise sequences, within sessions and throughout the entire experiment, showing a high degree of vocal plasticity. The F0 shift did not become persistent starting from a specific trial nor was induced by the natural growth of anatomical structures such as vocal folds [62]. Vocal production involving volitional modulations of acoustic parameters may highlight a rare ability in harbour seal pups. It has been previously shown that elaborated control over the vocal apparatus constitutes biophysical mechanisms for vocal learning. Thus, laryngeal plasticity and vocal flexibility may compose indirect evidence for vocal learning in harbour seals’ puppyhood [43, 62, 88].

F0 is a major feature shaping human singing and speech production. It is produced and controlled by the larynx. Frequency modulations may be physiologically more demanding to perform than temporal or amplitude modulations; in fact, several anatomical and dynamic features affect F0, such as length, tension, and rate of vibration of the vocal folds [62, 89]. Therefore, controlling F0 requires neuromuscular control over several anatomical structures whereas duration and amplitude of a sound are mostly controlled by modifications of exhalation.

In humans vocal learning requires control, mediated by the laryngeal motor cortex, over multiple phonatory structures linked to both the source and the filter [90]. Neurobiological studies, based on electrical stimulation and localised destructions, showed that the laryngeal motor cortex has a key role in volitional control of vocal fold movements [91, 92]. Direct cortico-bulbar connections have been suggested to be the main anatomical explanation for humans’ capacities of fine laryngeal control and vocal plasticity [93–95]. Few studies have demonstrated that those mammals incapable of vocal learning possess only *indirect* connections, which could explain their limited vocal plasticity [96, 97]. On the contrary, recent studies showed evidence that songbirds, that have demonstrated vocal learning abilities, might have analogous connectivity to humans [98]. To date, there is no evidence for direct cortico-bulbar projections in any mammalian species except for humans. Our behavioural results make harbour seals prime candidates among mammals to show direct anatomical connectivity between the laryngeal motor cortex and laryngeal motoneurons, as seen in humans [93, 99].

### Future work and conclusions

Parallel strands of research in mammalian bioacoustics suggest that harbour and grey seals are ideal model organisms to probe vocal learning in mammals [63, 99]. Mammalian vocal learning research can, in turn, shed light on the origins of human speech. Additional work on the level of control seals might exert over different parts of their phonatory apparatus can shed light on fine-grained mechanisms for vocal learning. Bats are also vocal learners and have been recently shown able to adjust various vocal parameters, independently from each other, in a setup comparable to ours [6]. Considering our relatively straightforward setup, we suggest that F0 modulation in response to noise could be a powerful cross-species test for vocal learning and plasticity, testing their association.

Further studies could investigate whether the vocal modulation of F0 in the presence of spectral masking is biologically relevant and actually perceived by conspecific harbour seals. Future work should also replicate the current experiment with increased noise levels to test the hypothesis that playbacks louder than ours could induce a stronger amplitude shift [20]. In addition, independent groups of seal pups could be exposed to different experiment noise playbacks. By varying the masking frequency band, its timing, and its intensity, one could test whether different frequencies could induce an upward vs. downward shift in F0. Comparing these conditions with masking frequencies that do not overlap with pups’ F0 or formants may trigger other types of compensatory adaptations instead of frequency shifts. Therefore, other experimental designs could spur the seals to perform amplitude shifts or temporal modifications, or instead even stronger frequency adaptations. As a complement to behavioural experiments, anatomical work could investigate the elastic properties of seal larynges to establish upper and lower anatomical boundaries for F0 production [100, 101]. Finally, neurobiological work should track purported direct cortico-laryngeal connections in seal pups, and compare them against closely related Caniformia not capable of F0 plasticity [90, 99].

To conclude, our data show plastic vocal behaviour in a neonate mammal, similar to that of humans and very few other adult mammals [6, 47]. As we learn more about vocal plasticity across species, we will be able to construct acoustic phylogenies of this trait in mammals. This will shed light not only on how environment and ancestry interact to deliver adaptable communication, but indirectly provide indirect inference on the evolution of speech and song in our own species.

## Acknowledgements

The authors would also like to thank all the volunteers, seal care members, and the veterinarians of the Zeehondencentrum Pieterburen that helped during the experiment and the rehabilitation process of the seal pups. We are grateful to Bart de Boer, Daria Valente, Marco Gamba, and Romain di Stasi for helpful discussions about audio analysis, statistics, or data interpretation. HR was funded by Ulla Tuominen Foundation.

## Author contributions

LTB, ASC and AR conceived the research and designed the experiment. LTB and AR collected the data, and LTB annotated it. YJ, HR, and AR extracted the acoustic features and performed the acoustic analyses. LTB, YJ, and AR performed the statistical analysis. All authors interpreted the results, drafted the article and revised it critically.

